# Nano-scale size holes in ER sheets provide an alternative to tubules for highly-curved membranes

**DOI:** 10.1101/191536

**Authors:** Lena K. Schroeder, Andrew E. S. Barentine, Sarah Schweighofer, David Baddeley, Joerg Bewersdorf, Shirin Bahmanyar

**Author notes:** These authors contributed equally to this work (Andrew E. S. Barentine, Sarah Schweighofer).

## Abstract

The endoplasmic reticulum (ER) is composed of interconnected membrane sheets and tubules. Super-resolution microscopy recently revealed densely packed, rapidly moving ER tubules, highlighting the importance of revisiting classical views of ER structure with high spatial resolution in living cells. Using live-cell Stimulated Emission Depletion (STED) microscopy, we show highly dynamic, subdiffraction-sized holes in ER sheets. Holes coexist with uniform sheet regions and are distinct from tubular ER structures. The curvature-stabilizing reticulon protein Rtn4 localizes to these holes and the ER luminal tether Climp63 controls their diameter and mobility. Analytical modeling demonstrates that holes in ER sheets can serve as reservoirs for curvature-stabilizing proteins to support ER tubule extension and retraction, thus providing an explanation for how the ER locally alters its morphology on fast time-scales.

**One Sentence Summary:** Dynamic nano-scale sized holes are prominent features of ER sheets that serve as reservoirs for curvature-stabilizing proteins to support ER tubule extension and retraction.

## Main Text

The endoplasmic reticulum (ER) is the largest membrane-bound organelle in eukaryotic cells. ER membranes are spread throughout the cytoplasm to perform essential functions in protein and lipid synthesis as well as calcium signaling. The continuous membranes of the ER extend from the nuclear envelope (NE) as stacks of ribosome-studded sheets and transition into an interconnected network of tubules and sheets at the cell periphery (*1*, *2*). Following the description of ER structure from electron micrographs over 60 years ago (*3*), fluorescence microscopy of living cells revealed that peripheral ER membranes are highly dynamic and constantly reorganize to facilitate the various functions of the ER (*1*, *2*, *4*). Further highlighting the importance of using advanced imaging techniques to gain a comprehensive view of ER morphology, recent work from Nixon-Abell et al. used structured illumination microscopy (∼100 nm resolution) to identify “ER matrices,” dense and highly dynamic ER tubule networks that appear to be sheets by conventional microscopy (*5*). This finding challenges the dogma that the peripheral ER consists of two distinct morphologies: flat sheets and curved tubules (*1*, *2*, *6*).

The high membrane curvature of ER tubules and rims of sheets require the ER-specific wedge-shaped Reticulon and DP1/Yop1 family of membrane inserted proteins, which are excluded from flat membrane sheet regions (*7*). The integral membrane protein Climp63 maintains the luminal space of ER sheets by forming bridges between parallel membrane sheets through its coiled-coil domain (*8*). While Climp63 overexpression induces ER sheets, the formation of ER sheets does not depend on Climp63. Instead, it is proposed that the abundance of Reticulon and DP1/Yop1 proteins relative to the amount of bilayer lipids determines the ratio of ER sheets to tubules (*8*).

To clarify discrepancies between electron microscopy (EM) and live-cell imaging about the presence of ER sheets at the cell periphery, we used Stimulated Emission Depletion (STED) microscopy, a super-resolution technique that allows fluorescence imaging of living cells at 50 nm resolution (*9*).

We imaged the periphery of live COS-7 cells expressing the genetically encoded fusion protein ss-SNAP-KDEL (*10*), which exclusively localizes to the ER lumen and can be labeled with organic dyes compatible with STED imaging (*11*) (Fig. 1A). Acquiring STED and confocal images of the same region within the cell periphery revealed ER structures that were not detected by conventional microscopy approaches (Fig. 1B, C). In regions that appeared to be ER sheets by confocal microscopy, STED microscopy revealed sheets containing distinct holes directly adjacent to uniform regions within the same sheet (Fig. 1C, right panels). Sheets containing holes were distinct from clusters of tubules, which also appeared as sheets by conventional microscopy (Fig. 1B, right panels). Holes in ER sheets were also observed in U-2 OS cells (Fig. 1D, E), with luminal and membrane ER markers (Fig. 1, fig. S1A), and in 3D image stacks of ER sheets near the nucleus (fig. S1B, C). Holes of similar diameter could also be observed in the NE, where nuclear pore complexes (NPCs) reside (fig. S1B-F). Due to their sub-diffraction limited size, we termed holes in ER sheets “nanoholes.”

**Fig. 1.**
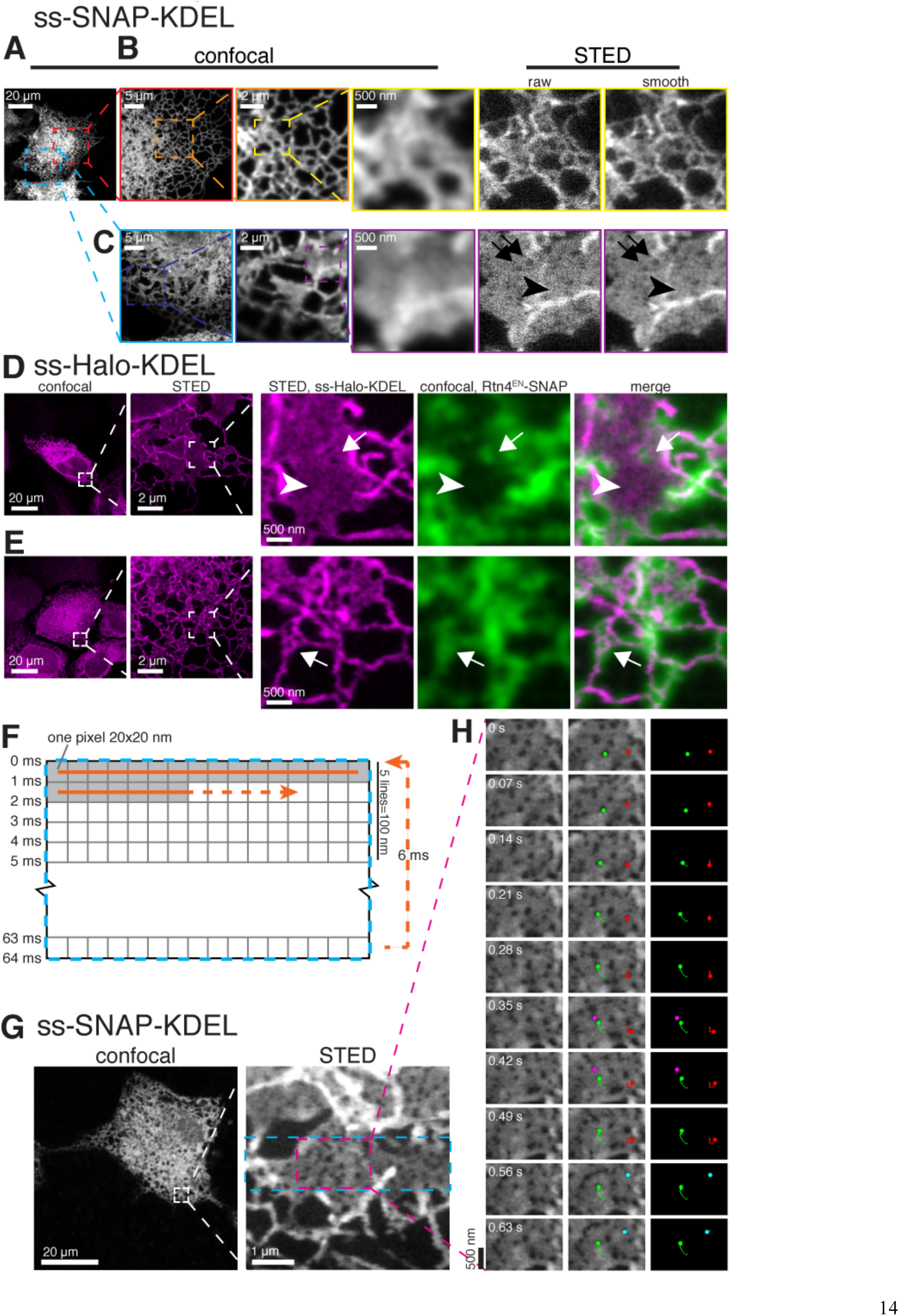
Nano-scale sized holes coexist with uniform regions in ER sheets. (**A-C**) Confocal and STED images of ss-SNAP-KDEL in living COS-7 cells. Arrowhead in *(C)* shows uniform sheet region near region with holes (arrows). Boxed regions in subsequent magnified live images may be slightly shifted because of ER movements. (**D-E**) Confocal and STED images of ss-Halo-KDEL in living U-2 OS cell (*magenta)*. Line sequential confocal images of endogenously tagged Rtn4-SNAP^EN^ (*green*). Arrowhead in *(D)* shows uniform ER sheet devoid of Rtn4-SNAP^EN^ and Rtn4-SNAP^EN^ localization to holes (arrow; referred to as “nanoholes”). Rtn4-SNAP^EN^ localizes throughout tubular network in *(E).* (**F**) STED image acquisition schematic. (**G**) Confocal image and magnified STED image of ss-SNAP-KDEL in living COS-7 cell. Blue rectangular region in magnified image outlines the 4.8 μm X 1.2 μm region imaged live with smaller magenta boxed region of the movie shown in *(H)*. (**H**) Select STED images from time-lapse series paired with images pseudocolored to highlight manually tracked nanoholes during time course of movie. Holes represented by green and red are persistent, by magenta appear and disappear, and by cyan appear. Time shown in seconds.

To determine if curvature-stabilizing reticulon proteins localize to curved membranes generated by nanoholes in ER sheets, we used CRISPR/Cas9 technology to introduce a SNAP tag at the endogenous locus of Reticulon 4 in U-2 OS cells (Rtn4-SNAP). Confocal imaging of Rtn4-SNAP in living cells revealed its colocalization to a subset of nanoholes in ER sheets that were identified by STED imaging of the luminal marker ss-Halo-KDEL (Fig. 1D, arrow). Rtn4-SNAP localizes to nanoholes that juxtapose uniform sheet areas completely devoid of Rtn4-SNAP (Fig. 1D, arrowhead). Rtn4-SNAP localization to nanoholes in flat sheets was distinct from membrane structures resembling clustered tubules that contain Rtn4-SNAP throughout (Fig. 1E). The specific localization of Rtn4 to nanoholes, but not to adjacent flat membrane sheet regions, was confirmed using two-color STED imaging of immunolabeled endogenous Rtn4 (fig. S2). We conclude that Rtn4 associates with nanoholes in otherwise flat ER sheets.

To determine whether nanoholes are stable or dynamic features of ER sheets, we took advantage of the rapid scan speed of the STED microscope’s resonant scanner in bidirectional scan mode (16,000 scanned lines per second) to capture 4.8 μm x 100 nm regions in 5 ms (Fig. 1F-H, movie S1). By combining the high spatial resolution provided by STED with excellent local temporal resolution, this approach was comparable to imaging of similar regions in the ER by structured illumination (fig. S1G, movie S2), but provided the superior resolution needed to clearly detect nanoholes (Fig. 1G, H, movie S1). This method shows that nanoholes are extremely mobile in the plane of ER sheets (Fig. 1H). Tracking individual holes (n = 24), we found that only 45% of nanoholes persisted during the time course of the movie. Surprisingly, 55% of tracked nanoholes appeared, disappeared or both during the time course of the movie (Fig. 1H, movie S1). These data suggest that the majority of nanoholes are transient or change between diameters above and below our resolving power on sub-second time scales.

To accurately characterize nanohole architecture in the context of other ER structures, we measured the improved resolution provided by STED microscopy. Rather than relying on measuring bead samples that are not directly comparable to our live-cell imaging experiments, we developed a method to determine the resolution of STED images of living cells from the image itself (*12*). Our measurements determined a STED xy resolution of 49.9 ± 4.4 nm (mean ± SD, Fig. 2A, full width at half maximum (FWHM) of the point-spread function (PSF)), about 5-times better than the diffraction limit, which facilitates the visualization of nanoholes in ER sheets and explains why nanoholes were not observed by conventional or structured illumination microscopy. Accounting for both the expected labeling of the structure, and the PSF of the microscope, we accurately determined the diameter of ER tubules in living cells (Fig. 2B-E, fig. S3) (*12*). We found that membrane-labeled ER tubules in live COS-7 cells are 101 ± 17 nm in diameter (mean ± SD, Fig. 2E), which is in agreement with prior reports that quantified ER tubule diameters from EM data (*13*, *14*). Using the same analysis on ER tubule data with luminal labeling yielded consistent diameters (107 ± 26 nm for ss-SNAP-KDEL and 115 ± 23 nm for ss-Halo-KDEL, mean ± SD, fig. S3).

**Fig. 2.**
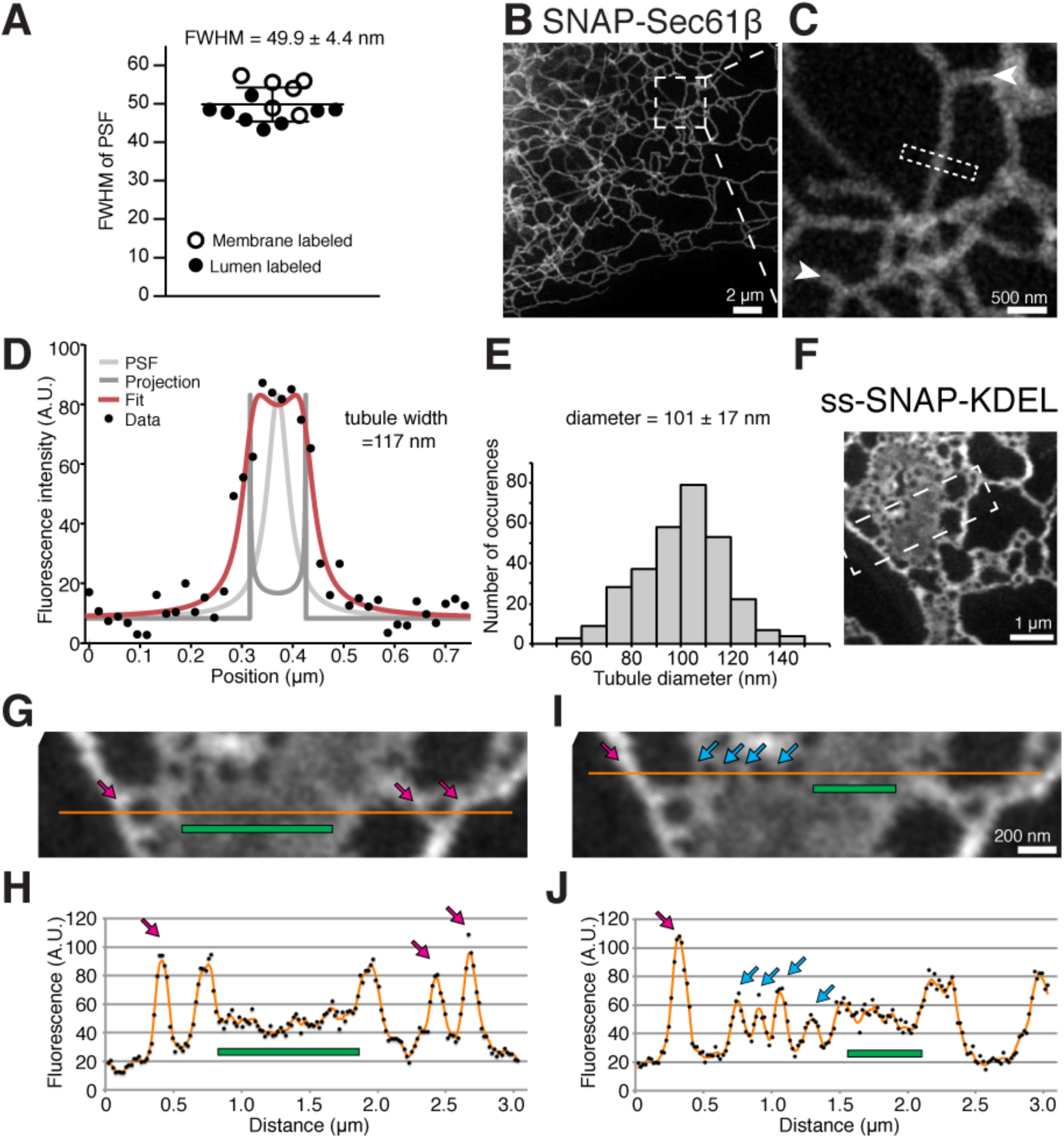
Analysis of tubules, sheets and nanoholes with STED resolution. (**A**) Plot of calculated PSF FWHM of STED microscope (N=4 imaging days; resolution 49.9 ± 4.4 nm, mean ± SD). (**B, C**) STED images of SNAP-Sec61β in living COS-7 cell. Arrowheads mark ER lumen. Dashed rectangle in *(C)* represents region used to generate fluorescence intensity line profile shown in *(D)*. (**D**) Plot represents fluorescence intensities along a line profile drawn perpendicular to an ER tubule (black dots; rectangular region in *(C)*) overlaid with the best fit (red line) of a projected line profile of an idealized surface-labeled cylinder (dark grey) convolved with a Lorentzian PSF (light grey). The best fit line has a 117 nm tubule diameter. (**E**) Histogram of SNAP-Sec61β labeled ER tubule diameters (n = 300 tubule profiles; N = 3 imaging days; 101 ± 17 nm, mean ± SD). (**F**) STED image of ss-SNAP-KDEL in living COS-7 cell. Boxed region magnified in (*G*, *I*). **(G-J)** Orange line in *(G, I)* represents five-pixel wide fluorescence intensity line profile shown in *(H, J)*. Magenta arrows mark ER tubules, green bars mark uniform ER sheets, and cyan arrows mark ER regions flanking nanoholes. Line profiles in *(H, J)* are from raw image (black dots) and image smoothed with uniform 3 x 3 filter (orange line).

To also estimate the thickness of ER sheets in living cells, we analyzed the intensity of the luminal ER marker ss-SNAP-KDEL. Assuming ss-SNAP-KDEL is evenly distributed throughout the ER lumen, the signal intensity of ss-SNAP-KDEL is roughly proportional to the thickness of the luminal space because the z-resolution of the STED microscope over which the signal is accumulated in the depth direction is much larger than the thickness of a single ER tubule or sheet (∼600 nm z-resolution versus 50-150 nm ER lumen thickness in Fig. 2E) (*15*). Line profiles drawn across images of ER sheets and tubules (Fig. 2F-J) show clear differences in the ss-SNAP-KDEL signal intensity between areas identifiable as flat sheets and tubules (Fig. 2H, J). The observed lower signal in ER sheets versus tubules (Fig. 2H) suggests that the thickness of ER sheets in living cells is approximately 30-50 nm thick (fig. S4), which is consistent with previous EM studies on fixed samples (*8*).

The signal intensity of ss-SNAP-KDEL in regions between nanoholes is similar to the signal intensity of uniform sheets and clearly lower than the signal intensity of tubules (Fig. 2I, J). These data, combined with high spatial resolution that resolves gaps larger than a few tens of nanometers and temporal resolution that shows the persistence of nanoholes, provide further evidence that we are detecting nanoholes in ER sheets rather than gaps in a matrix of highly dynamic tubules.

To gain a more comprehensive understanding of cellular factors that affect nanohole architecture, we developed a semi-automated algorithm using watershed transformation that detects hole borders to measure nanohole size and shape in an unbiased manner (Fig. 3A-E). As a reference, we applied the algorithm to analyze holes in the NE. The measured symmetric shape of NE holes (Fig. 3D, fig S5), tight distribution of NE hole diameter (Fig. 3E, fig S5), and their uniform spatial distribution (fig. S1D, F) supports the interpretation that they indeed represent regions at which NPCs are inserted. The shape and size of nanoholes in ER sheets varied greatly compared to the tight distribution of the shape and size of NE holes (Fig. 3A, D, E). This difference suggests that the structure of nanoholes in ER sheets is regulated by a mechanism distinct from that responsible for forming and stabilizing NPCs.

**Fig. 3.**
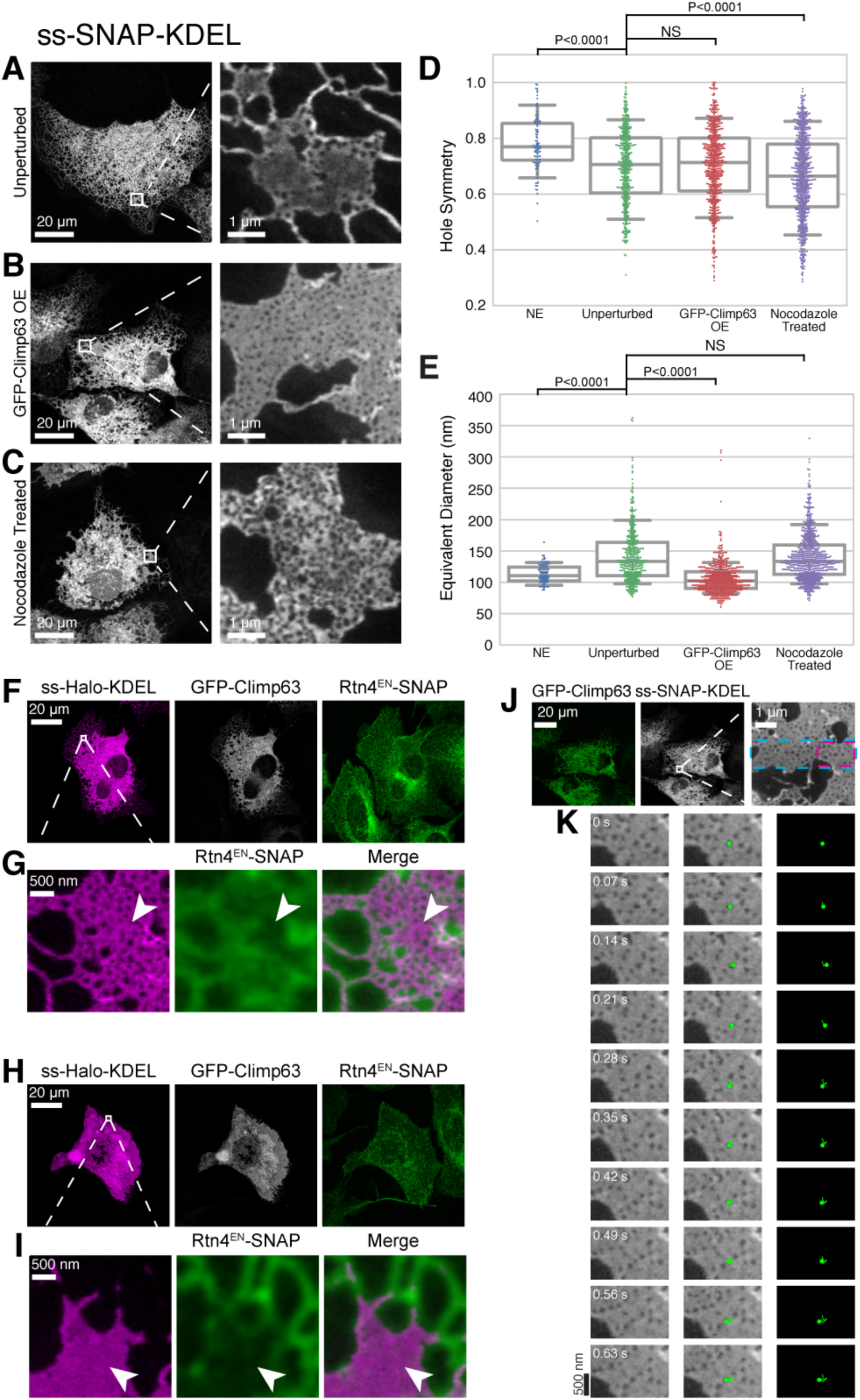
Architecture of nanoholes in ER sheets. (**A-C**) Confocal images of ss-SNAP-KDEL in living COS-7 cells (*left*) and STED image of magnified boxed region (*right*) in indicated conditions. (**D, E**) Plot of nanohole symmetry and nanohole equivalent diameter in indicated conditions. Kruskal-Wallis and Dunn’s test determined p values. NS = not significant. (NE, n= 97 holes, N= 3 cells; Unperturbed, n= 424 holes, N= 2 cells; GFP-Climp63, n= 601 holes, N= 3 cells; and Nocodazole treatment, n= 713 holes, N= 4 cells) (**F**) Confocal images of U-2 OS cell expressing Rtn4-SNAP^EN^ (*green*), ss-Halo-KDEL (*magenta*), and GFP-Climp63 (*grayscale*). (**G**) STED image of ss-Halo-KDEL from boxed region in *(F)* (Rtn4-SNAP^EN^ taken at confocal resolution). Arrowhead marks uniform membrane regions. **(H)** Confocal images as described in *(F)*. **(I)** STED image of ss-Halo-KDEL from boxed region in *(H)*. Arrowhead marks Rtn4-SNAP^EN^ displaced from ER sheets. (**J**) Fluorescence confocal image of cells in (*B*) expressing GFP-Climp63 (*green*) and ss-SNAP-KDEL (*grayscale*). Dashed blue line in right panel image outlines the 4.8 μm X 1.2 μm region imaged over time. The region outlined by the dashed magenta boxed is shown in *(K)*. (**K**) Select STED images of ss-SNAP-KDEL paired with pseudocolored image and image overlay (third column) to highlight a manually tracked nanohole that persist. Time shown in seconds.

Next, we tested how changing the abundance of sheets relative to tubules affects nanohole size and shape by overexpressing the intraluminal spacer Climp63 (*8*). Our analysis showed that Climp63 overexpression in COS-7 cells constricts nanoholes to a diameter close to that of holes in the NE (median values 102 nm and 111 nm, respectively, vs. 133 nm in unperturbed cells) (Fig. 3A, B, D, and E). In comparison to cells expressing Climp63 at endogenous levels, these sheets also featured remarkably uniform ss-SNAP-KDEL signal in areas between nanoholes indicating even ER sheet thickness (Fig. 3B; compare to Fig. 3A or Fig. 2F). Endogenously tagged Rtn4-SNAP localized to nanoholes in Climp63-induced sheets and was absent from flat membrane regions (Fig. 3F, G). High overexpression of Climp63 generated large uniform sheets that did not contain nanoholes and displaced Rtn4-SNAP from nanoholes to the few remaining tubules (Fig. 3H, I). The observation that overexpression of Climp 63 constricts nanohole size and Rtn4 localizes to nanoholes is consistent with a role for Climp63 in tethering parallel sheets across the lumen, which stabilizes flat membranes, and the preference of Rtn4 for curved edges (*8*).

To test if generating ER sheets by a different method also affected nanohole architecture, we destabilized microtubules by treating cells with 33 μM nocodazole, which causes the peripheral ER to coalesce into sheets (*16*). STED microscopy revealed ER structures generated by microtubule depolymerization that resembled closely packed tubular networks rather than sheets (Fig. 3C), and contained endogenous Rtn4 throughout (fig. S6). The increased spread in the distribution of hole symmetry (Fig. 3D) reflects the change in ER morphology caused by nocodazole treatment (Fig. 3A, C). Thus, both microtubule depolymerization and low Climp63 overexpression do not support expanded ER tubular networks, yet they result in different ER morphologies, which are distinct from uniform sheets.

We further reasoned that increasing the abundance of Climp63, which has been reported to multimerize into clusters with low mobility (*17*), may corral nanoholes in ER sheets and limit their ability to travel in the plane of the sheet. We were able to manually track a greater number of nanoholes in Climp63-induced sheets than in untreated cells (Fig. 3J, K compared to Fig. 1G, H), and a greater percentage of nanoholes persisted over the imaging time course (80%, n=59 holes, Fig. 3K). These data indicate that the dynamics of nanoholes in ER sheets are strongly influenced by Climp63. The increased persistence of nanoholes in ER sheets induced by Climp63 overexpression provides further evidence that nanoholes in ER sheets are distinct from highly dynamic tubules that form matrices in the cell periphery (*5*).

We next addressed the significance of Rtn4 localization to nanoholes in ER sheets. We hypothesized that nanoholes in ER sheets may act as a readily available reservoir for curvature-stabilizing proteins, such as Rtn4, that facilitate tubule formationand thereby support morphological changes to the ER more quickly than through protein or lipid synthesis or interorganelle transfer. To investigate how the amount of curved membrane changes as a function of its morphology, we developed a simple geometric model of the ER consisting of a single disk-shaped sheet, with an adjustable number of nanoholes (100 nm inner diameter) and a 100-nm diameter ER tubule of variable length extending from it (Fig. 4A). We assumed a 50-nm thick ER sheet, which is consistent with our thickness estimates in living cells (fig S4; modeling with other sheet thicknesses in fig S7) as well as previous reports in fixed cells (*8*). We exclude lipid synthesis or interorganelle transfer from our model by assuming that the total surface area of the modeled ER is constant and any change in surface area resulting from tubule or nanohole addition is compensated by an adjustment of the sheet disk diameter, which is initially 5 μm. Using this geometric model, we determined that ten nanoholes store the same curved membrane area as ∼1 μm of ER tubule (Fig. 4B, C). Notably, the curved surface area of a hole in a 50-nm thick sheet has a higher degree of curvature and can hold more wedge-shaped curvature-stabilizing proteins than a 100-nm thick tubule of equal surface area (see schematic in fig. S8A, B). Overall, we show that a modest number of nanoholes contain sufficient curvature to support tubule extension or retraction, without changes in the total number of curvature-stabilizing proteins or lipids.

**Fig. 4.**
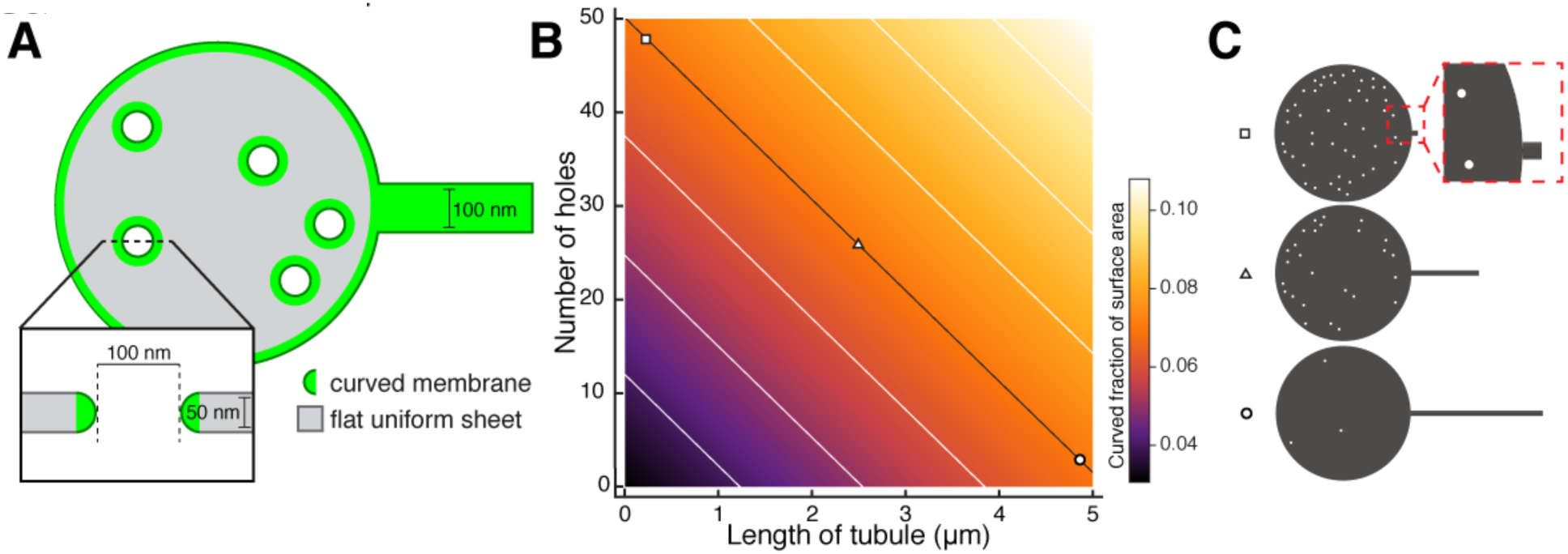
Analytic model of curved edges of nanoholes serving as a reservoir for tubule extension and retraction. (**A**) Schematic of geometries used to model the ER: the sheet is assumed to be a disk with half-toroid edges, empty cylinders with half-toroid edges represent nanoholes, and a cylinder describes the tubule. A cross section of a nanohole is shown. (**B**). Heatmap of curved fraction of surface area as number of holes and tubule length are varied with the total surface area held constant. The essentially linear isoclines, where curved surface area is constant despite changes in tubule or hole number, are shown in white and black, with three points marked by shapes which are graphically represented in *(C)*. (**C**) Three points on the black isocline in *(B)*, coordinates marked with square, triangle, and circle are drawn to scale with the appropriate number of holes and length of tubule.

In summary, these results support our hypothesis that nanoholes in ER sheets provide a mechanism for forming ER tubules without requiring net change in lipids or proteins. Thus, in addition to the abundance of curvature stabilizing proteins and bilayer lipids in dictating ER morphology (*8*), nanoholes may provide a way to alter local ER morphology and dynamics on rapid time scales. The high mobility of nanoholes, which is strongly influenced by the expression level of the sheet stabilizer Climp63, could allow for their rapid diffusion to the sheet edge so that reticulons stored in nanoholes can easily assemble onto growing ER tubules. Altering local ER morphology with nanoholes may be important to ER functions that are specific to the cell periphery, especially at locations far away from the site of transcription or translation, such as at remote neuronal processes.

The fact that nanoholes have sheet-like as well as tubule-like morphological features allows for a diverse range of mechanisms to explain how they are created and resolved. In addition to machinery that can facilitate sheet fusion or fission (*18*, *19*), processes that enable ER tubule fusion and fission may control nanohole creation or removal within sheets (*6*) (see schematic in fig. S8C, D).

In conclusion, we experimentally demonstrate the existence of nano-scale sized fenestrations in ER sheets of living cells as a morphology clearly distinct from uniform sheets and ER matrices (*5*). Additionally, we find that ER sheets are present at the cell periphery, as has been reported by others using conventional and EM approaches (*6, 8, 20, 21*). ER morphology is linked to human disease (*1*), and thus the prominence of nanoholes as a membrane feature of the ER in living cells provides a more comprehensive view of ER structure that may contribute to our future understanding of the relationship between ER structure and disease.

## Acknowledgments

This work is supported in part by NIH grant S10 OD020142 for imaging resources (for the Leica TCS SP8 STED 3X microscope), the G. Harold & Leila Y. Mathers Foundation, the Wellcome Trust (095927/A/11/Z, 203285/B/16/Z), and the Yale Diabetes Research Center (NIH P30 DK045735). A. E. S. B. acknowledges support by an NIH training grant (T32 GM008283). S. S. acknowledges support by the Austrian Marshall Plan Foundation. J. B. discloses significant financial interest in Bruker Corp. and Hamamatsu Photonics. We thank BioVision Technologies for the use of the VT-iSIM.

## Supplementary Materials

Materials and Methods

Figures S1-S8

Movies S1-S3

References (22-26)

Auxiliary Mathematica file (.pdf and .nb files, script of analytical modeling)

## Reference Material

Barentine *et al.* 2017 (reference 12, describing method used for Fig. 2)

